# Repeated shrimp allergen exposure drives 5-lipoxygenase-dependent avoidance and selective gut-brain activation

**DOI:** 10.64898/2026.09.09.750437

**Authors:** Camila Mattos Andrade, Noemi de Souza Pinto, Bruna Genisa Costa Lima, Ana Cristina Roginski, Ítalo da Silva Gonçalves, Raphael Chagas Silva, Letícia Marques Pilger, Amanda Maria Lago de Castro Leal, Gabriela Duarte da Silva, Natalia Machado Tavares, Tatiani Uceli Maioli, Ana Maria Caetano Faria, Cláudia Ida Brodskyn, Esther Borges Florsheim

## Abstract

Peripheral immune processes can shape animal behavior, yet how noninfectious inflammatory reactions affect neural activity and behavioral outputs remains poorly understood. We developed an optimized murine model of shrimp allergy using whole shrimp extract to examine how a complex dietary allergen elicits integrated immune, neural, and behavioral responses. Sensitized mice received repeated oral shrimp challenges and were assessed for allergic pathology, food preference, affective-like behaviors, and neuronal activation in the brain. Repeated exposure increased total IgE and shrimp-specific IgG1, induced mast cell activation, accelerated gastrointestinal transit, caused mild hypothermia consistent with oral anaphylaxis, and increased intestinal length. Shrimp-sensitized mice did not avoid shrimp solution after sensitization alone. Instead, avoidance emerged only after repeated oral challenges and strengthened over time. This delayed aversion occurred without detectable changes in locomotor activity or measures of anxiety-like or depressive-like behavior at the time points tested. Repeated shrimp exposure increased cFOS expression in the area postrema, nucleus of the tractus solitarius, central amygdala, and paraventricular nucleus of the thalamus, implicating brainstem and limbic-thalamic pathways involved in visceral sensing and aversion. Pharmacological inhibition of 5-lipoxygenase partially reversed avoidance and reduced circulating mast cell protease-1 in allergic mice. These findings establish a robust whole-shrimp allergy model and show that a complex food allergen engages gut-brain pathways to promote 5-lipoxygenase-dependent avoidance. The delayed and selective nature of this response supports immune-mediated food aversion as a shared output of food allergy while suggesting that its kinetics and neural recruitment vary with allergen identity and inflammatory context.

## INTRODUCTION

To survive in chemically complex environments, animals must detect harmful substances and coordinate physiological and behavioral defenses through immune, neural, and endocrine pathways. Infection-induced sickness behaviors are an example of this integration as inflammatory signals decrease appetite and locomotion, promote fatigue, social withdrawal, and avoidance of contaminated resources^1,2^. It is thought that these responses are not merely passive consequences of illness, but regulated defense programs that can promote recovery and survival^2–5^. In contrast, much less is known about how distinct inflammatory responses arising in noninfectious settings, including allergic type 2 immunity, shape brain function and behavior. Although type 2 immunity contributes to host defense against parasites, venoms, and toxins^6–8^, it remains poorly understood whether allergic inflammation induces classical sickness-like states or distinct behavioral programs.

Food allergy provides a relevant model to define how type 2 inflammation in the gastrointestinal tract connects with the nervous system to drive behavioral outputs^9^. In clinical settings, food allergy has been associated with neuropsychiatric symptoms, including anxiety, suggesting that hypersensitivity can extend beyond peripheral reactions^10–12^. Experimentally, subclinical models of cow’s milk allergy have shown that oral sensitization to whey proteins can alter behavior, neuroinflammatory responses, and central histaminergic signaling^13–15^. In parallel, ovalbumin (OVA)-induced food allergy models have demonstrated that allergic sensitization can promote antigen-specific avoidance through IgE-, mast cell-, and leukotriene-dependent mechanisms^16–20^. This avoidance is associated with activation of visceral-sensing and aversion-related brain regions, including the nucleus of the tractus solitarius (NTS), external lateral parabrachial nucleus (elPBN), and central amygdala (CeA)^18,20,21^. Together, these studies indicate that allergic inflammation engages gut–brain pathways to shape behavior.

Whether these neural and behavioral responses are conserved features of food allergy or depend on allergen identity and exposure context remains unclear. The few mechanistic studies of allergen-induced avoidance have relied on OVA, a well-defined and major egg allergen that provides a tractable single-protein system. However, hypersensitive individuals typically encounter allergens within complex foods, where multiple allergenic and non-allergenic components may influence sensitization, digestion, innate immune activation, and inflammatory responses^22–25^. Food allergens also differ in biochemical composition, abundance, structural stability, and digestion resistance, all of which may alter the magnitude, timing, and quality of gut inflammation and, therefore, brain responses. Hence expanding beyond single-protein allergen models is important for defining which features of allergy-induced behavior are broadly conserved and which are shaped by allergen identity and dietary exposure context.

In this study, we established an optimized and more robust murine model of shrimp allergy using whole shrimp extract to test whether a complex dietary allergen promotes immune-mediated behavioral and neural responses. Shrimp is also clinically pertinent, as shellfish allergy is common in adults and is a major cause of food-induced anaphylaxis^26,27^. Sensitized mice developed hallmark features of food allergy, including elevated total IgE, shrimp-specific IgG1, mast cell activation, mild oral anaphylaxis, and accelerated gastrointestinal motility. Unlike OVA-induced models, shrimp-induced avoidance emerged only after repeated oral allergen exposure and coincided with the development of allergic inflammation. This avoidance required 5-lipoxygenase activity and was accompanied by activation of aversion-and visceral-sensing brain regions, including the NTS, CeA, area postrema, and paraventricular nucleus of the thalamus. Notably, shrimp allergy did not produce persistent locomotor changes or a consistent affective-like phenotype at the time points tested, suggesting that the sustained behavioral response was primarily directed toward the allergen rather than a generalized sickness-like state. Together, these findings support immune-dependent dietary avoidance as a shared neuroimmune defense strategy across distinct food allergy models, while showing that allergen identity and exposure context shape its timing, inflammatory threshold, and neural recruitment.

## RESULTS

### Repeated oral exposure to shrimp induces key hallmarks of food allergy in sensitized mice

Shrimp allergy is clinically important but remains difficult to study mechanistically because of the limited availability of experimental models that recapitulate major features of the disease. We previously established a mouse model of shrimp allergy that induced mild allergic responses^28^. Here, we optimized this model to generate more robust allergic pathology and to investigate the immune, neural, and behavioral consequences of repeated oral exposure to dietary shrimp.

Mice were sensitized with shrimp extract and alum on days 0 and 14, whereas control animals received PBS plus alum. Beginning on day 21, mice were challenged intragastrically with shrimp extract six times (Figure 1A). Shrimp-sensitized mice developed elevated total IgE levels, which further increased after repeated oral challenges compared with controls (Figure 1B). Similarly, sensitized mice showed increased shrimp-specific IgG1 after the final sensitization on day 20, and these levels were further boosted after oral challenges on day 37 (Figure 1C). Repeated oral shrimp exposure also increased shrimp-specific IgG1 in control animals, indicating that oral exposure alone was sufficient to elicit a detectable IgG1 response.

**Figure 1.**
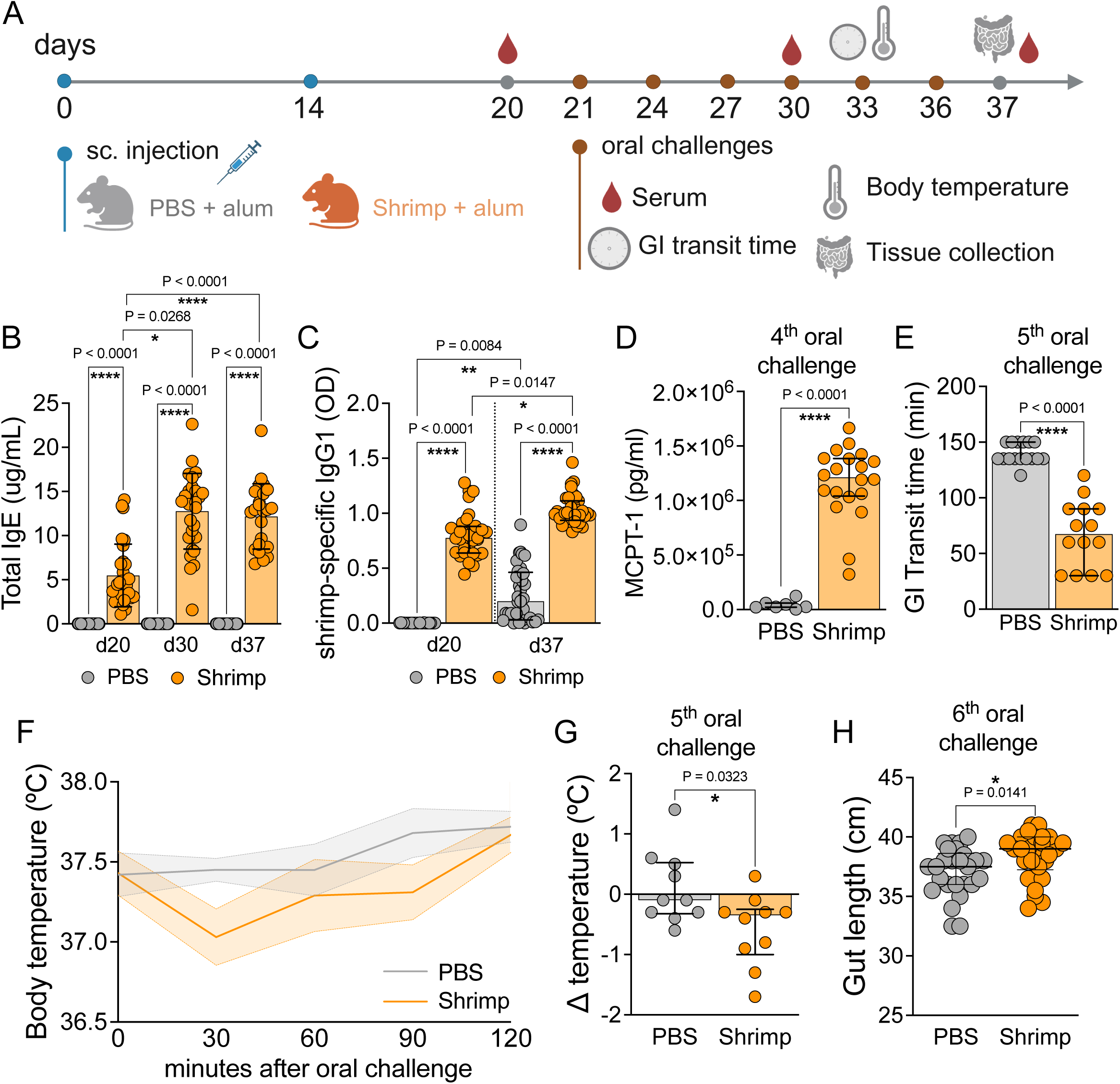
Chronic oral exposure to shrimp induces major hallmarks of food allergy in an experimental BALB/c mouse model. (**A**) Schematic protocol for allergic sensitization, oral challenges, and physiological read outs. (**B**) Total levels of serum IgE after allergic sensitization (day 20), after oral challenges (day 30 and day 37) in BALB/c mice sensitized with PBS or shrimp + alum (n= 20 control and 25 allergic per group). (**C**) Shrimp-specific levels of serum IgG1 after allergic sensitization (day 20) and oral challenges (day 37) in BALB/c mice sensitized with PBS or shrimp + alum (n= 20 control and 25 allergic per group). (**D**) MCPT-1 levels 1h after fourth oral challenge from control and allergic mice sera (n = 10 mice per group). (**E**) Gastrointestinal transit time was determined on the fifth oral challenge with shrimp using a red carmine assay (n = 18 mice per group). (**F**) Rectal temperature over time and (**G**) maximum temperature variation (Δ) after fifth oral challenge in PBS or shrimp + alum-sensitized mice (n = 10 mice per group). (**H**) Gut length was determined by direct measurement of the small intestine using ruler 24h after sixth oral challenge (n=28 mice per group). Graphs shown as median±interquartile range (**B, C, D, E, G, H**) and mean±s.e.m (**F**). *P≤0.05, **P≤0.01, ****P≤0.0001. Kruskal-Wallis with Dunn’s multiple comparisons test (**B, C**) and Two-tailed Mann–Whitney test (**C-H**). Each panel is representative of at least two independent experiments. (**A**) Created with BioRender.com.

Consistent with mast cell activation, shrimp-sensitized mice had increased serum levels of mast cell protease-1 (MCPT-1) one hour after the fourth oral challenge (Figure 1D). After the fifth challenge, sensitized mice also displayed accelerated gastrointestinal transit compared with controls, without overt diarrhea, indicating increased intestinal motility (Figure 1E). Repeated oral shrimp exposure further induced a mild decrease in body temperature in sensitized mice, consistent with oral anaphylaxis and an IgE-associated allergic reaction (Figures 1F–G). After six oral challenges, allergic mice also showed increased intestinal length, suggesting local inflammation and tissue remodeling (Figure 1H). Together, these findings indicate that repeated oral exposure to shrimp induces key immunological and physiological features of experimental food allergy in sensitized mice.

### Shrimp allergy promotes avoidance behavior only after repeated oral allergen exposure

Avoidance of food allergens has been characterized primarily in OVA-induced food allergy models, in which sensitized animals rapidly avoid the allergen after a single oral exposure^16,19,20^. To determine whether shrimp allergens induce a similar behavioral response, we assessed shrimp preference in sensitized and control mice using a two-bottle choice assay.

After the final sensitization, mice were acclimated to two bottles containing water (Figure 2A). During acclimation, both groups showed a similar preference for the bottle positioned on the right, indicating a positional bias unrelated to allergic sensitization (Figure 2B). To control for this bias, the position of the shrimp-containing bottle was alternated across testing days. When one water bottle was replaced with a shrimp-containing solution, shrimp-sensitized mice did not show reduced preference for the shrimp solution during the 3-day testing period (Figures 2C–D), despite having higher total IgE levels than control mice (Figure 1B).

**Figure 2.**
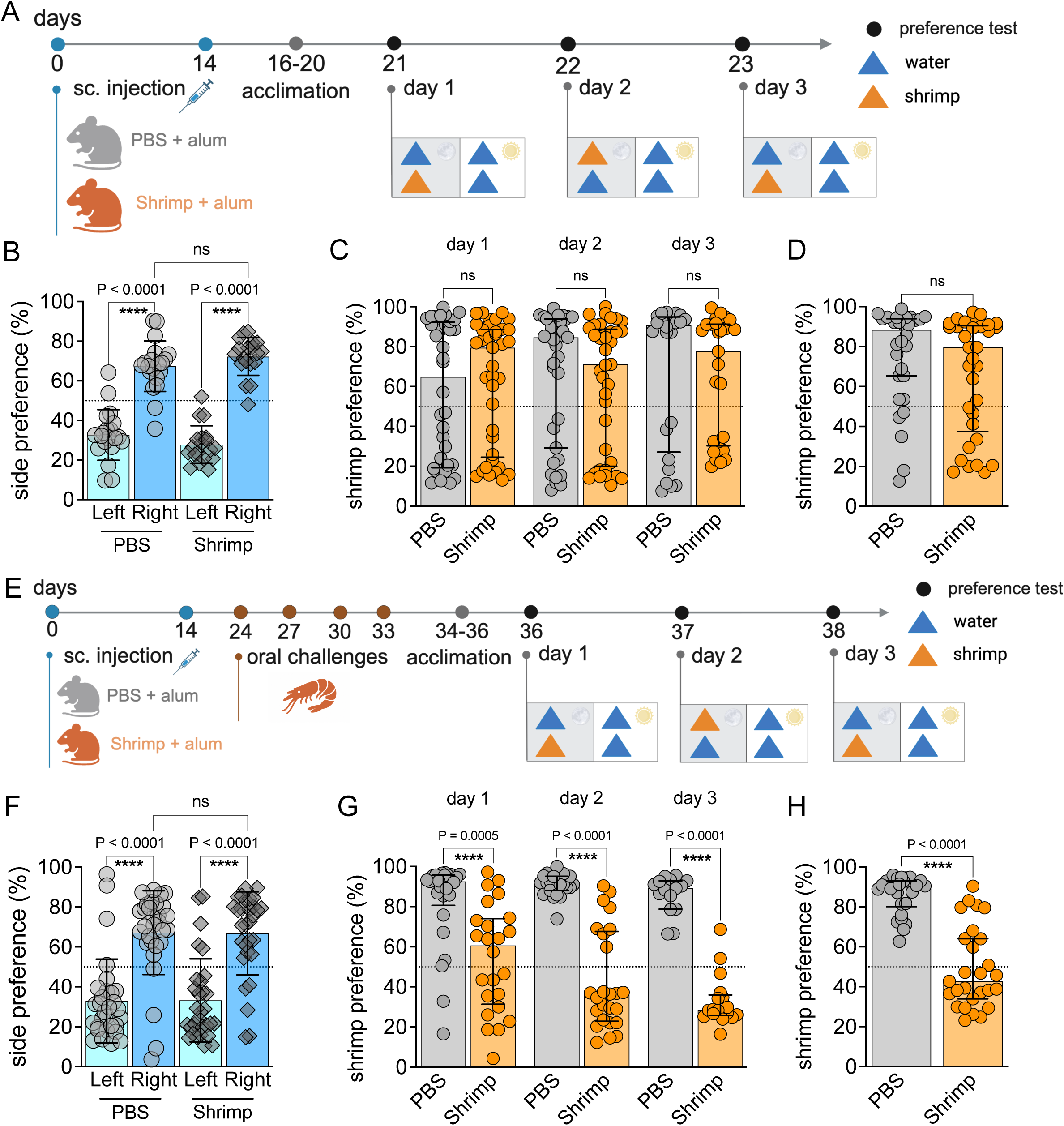
Shrimp allergy promotes avoidance behavior only after repeated oral exposure to allergen. (**A**) Schematic protocol for allergic sensitization and two-bottle preference test. (**B**) Mice were acclimated to two bottles of water for 3 days, and the side preference percentage was calculated. (**C**) Preference to shrimp solution from 3 days of preference test, consisting of one water bottle and one 0.5% shrimp bottle before the oral challenge, with bottle positions switched from day 2, in mice sensitized with PBS or shrimp + alum (n=28 mice per group). (**D**) Mean shrimp preference from the three days of test before oral challenges. (**E**) Schematic protocol for allergic sensitization, oral challenges, and two-bottle preference test. (**F**) Mice were acclimated to two bottles of water for 3 days, and the side preference percentage was calculated. (**G**) Shrimp preference after four oral challenges, with bottle positions switched from day 2, in mice sensitized with PBS or shrimp+alum (n=36 mice per group). (**H**) Mean shrimp preference from the three test days after oral challenges. Graphs shown mean±s.e.m. \**P*≤0.05, \*\**P*≤0.01, \*\*\*\**P*≤0.0001. (**B**, **E**) One-way analysis of variance (ANOVA) with Tukey’s multiple-comparison test. (**C**, **F**) Two-tailed Mann–Whitney test. Each panel is representative of at least three independent experiments. (**A**, **D**) Created with BioRender.com.

We next asked whether repeated oral allergen exposure was required to induce avoidance. After multiple oral shrimp challenges, mice were tested again on day 34 (Figure 2E). During acclimation, mice again retained a right-side preference independent of the experimental group (Figure 2F). In contrast to the post-sensitization test, shrimp-sensitized mice showed reduced preference for shrimp-containing solution after repeated oral exposure (Figures 2G–H). This avoidance was evident from the first day of testing and strengthened over time, with most allergic mice avoiding the shrimp solution by day 3. These findings indicate that, unlike OVA-induced models, shrimp allergy promotes allergen-specific avoidance only after repeated oral allergen exposure, coinciding with the development of allergic inflammation and physiological symptoms.

### Shrimp allergy induces selective avoidance without a consistent affective-like or locomotor phenotype

Food allergy is clinically associated with increased anxiety and reduced quality of life, particularly in patients at risk of severe reactions or accidental allergen exposure^10,11^. Given the high anaphylactic potential and lifelong persistence of shrimp allergy, we asked whether repeated oral exposure to shrimp allergens would promote anxiety-like behavior or other affective behavioral changes. To test this, we assessed shrimp-sensitized and control mice using the open field test (OFT), elevated plus maze (EPM), and forced swim test (FST) (Figure 3A).

**Figure 3.**
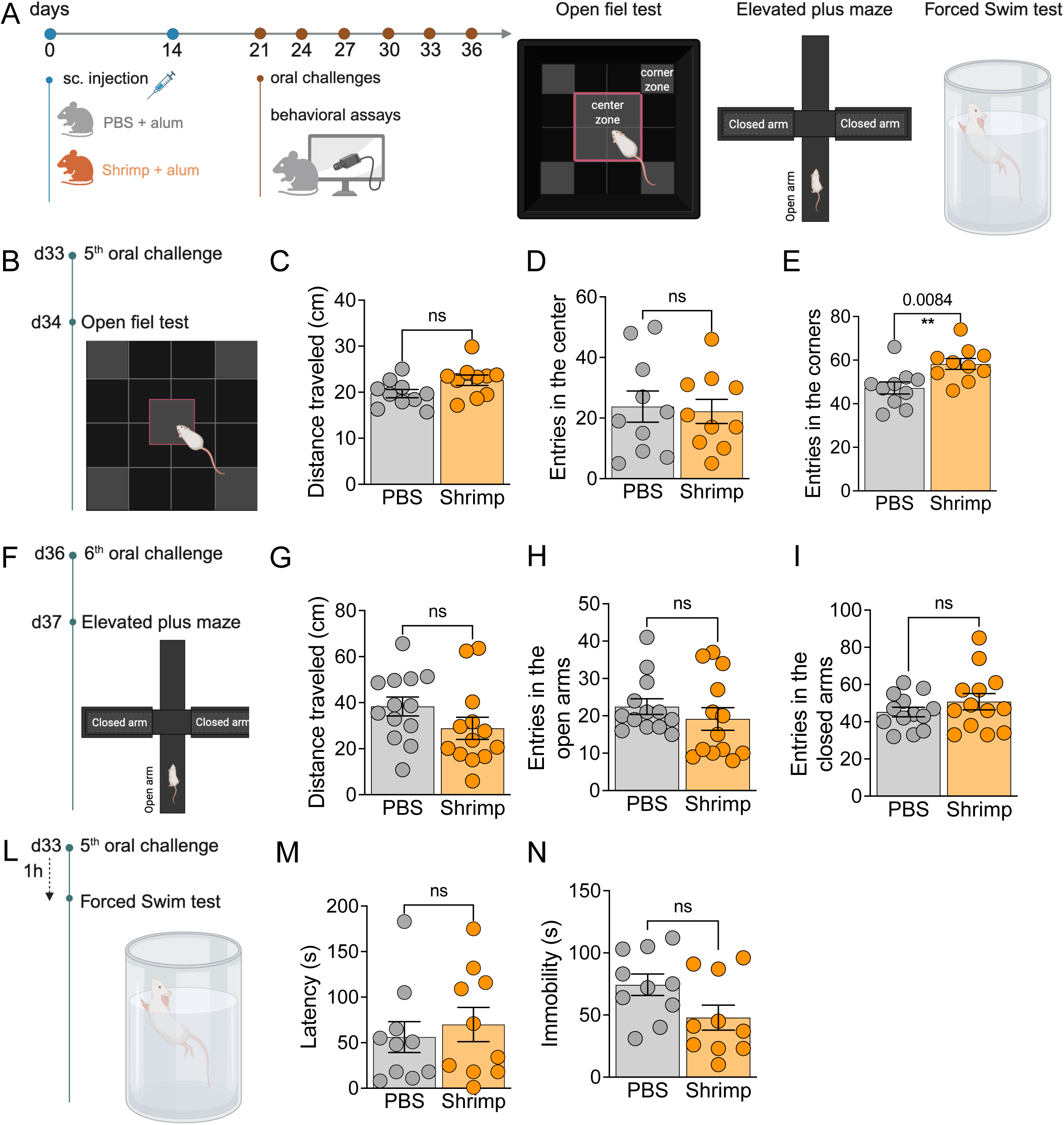
Shrimp allergy does not produce a consistent affective-like or locomotor phenotype at the time points tested. (**A**) Schematic protocol for allergic sensitization, oral challenges, and behavioral assays. (**B**) 24h after the 5th oral challenge, the open field test (OFT) was performed and recorded for 10 min per mouse. (**C**) Distance traveled, (**D**) entries in the center and (**E**) entries in the corners from OFT performed 24 hours after the 5th oral challenge in mice sensitized with PBS or shrimp + alum (n=15 mice per group). (**F**) 24h after the 6th oral challenge, the elevated plus maze (EPM) was performed and recorded 10 min per mouse. (**G**) Distance traveled, (**H**) entries in the open arms and (**I**) entries in the closed arms from EPM performed 24 hours after the 6th oral challenge in mice sensitized with PBS or shrimp + alum (n=15 mice per group). (**L**) 1 hour after the 5th oral challenge, the forced swim test (FST) was performed and recorded for 6 min per mouse. (**M**) Latency time and (**N**) immobility time from FST. Graphs shown mean±s.e.m.\*\**P*≤0.01. Two-tailed Unpaired T-test. Each panel is representative of at least two independent experiments. (**A**, **B**, **F**, **L**) Created with BioRender.com.

We first performed the OFT on day 22, 24 hours after a single oral shrimp challenge (Figure S1A). No differences were observed between control and shrimp-sensitized mice in time spent in the center or corners of the arena (Figures S1C-D). Sensitized mice showed a higher number of center entries (Figure S1E), but corner entries were unchanged (Figure S1F), and overall locomotor activity did not differ between groups (Figure S1B).

After repeated oral challenges, the OFT was performed again on day 34 (Figures 3B, S1G). Although both total locomotor activity and center entries remained similar between groups (Figure 3C-D), shrimp-sensitized mice showed increased corner entries compared with controls (Figures 3E). The amount of time spent in the center or corners was unchanged (Figures S1H-I). We then performed the EPM 24 hours after the sixth oral challenge (Figures 3F, S1L). Shrimp allergy did not alter locomotor activity, as measured by distance traveled during the assay (Figure 3G), the number of entries into the open or closed arms (Figures 3H-I), or the time spent in either arm (S1M-N).

To evaluate depressive-like behavior, we performed the FST one hour after the fifth oral shrimp challenge (Figure 3L). No differences were detected between groups in latency to immobility or total immobility time (Figures 3M–N). Although sensitized mice showed isolated changes in center or corner entries, these effects were not accompanied by consistent changes in center occupancy, EPM behavior, locomotor activity, or FST immobility. Thus, repeated shrimp exposure did not produce a consistent locomotor, anxiety-related, or immobility phenotype at the time points tested.

### Repeated oral shrimp exposure activates brain regions associated with visceral sensing and aversion

Aversive responses to noxious stimuli have been shown to engage brain regions such as the nucleus of the tractus solitarius (NTS), external lateral parabrachial nucleus (elPBN), and central amygdala (CeA)^18,20,29,30^. To determine whether repeated oral shrimp exposure activates similar brain regions, brains were collected 90 min after the fifth oral challenge, and neuronal activation was assessed by cFOS staining (Figure 4A).

**Figure 4.**
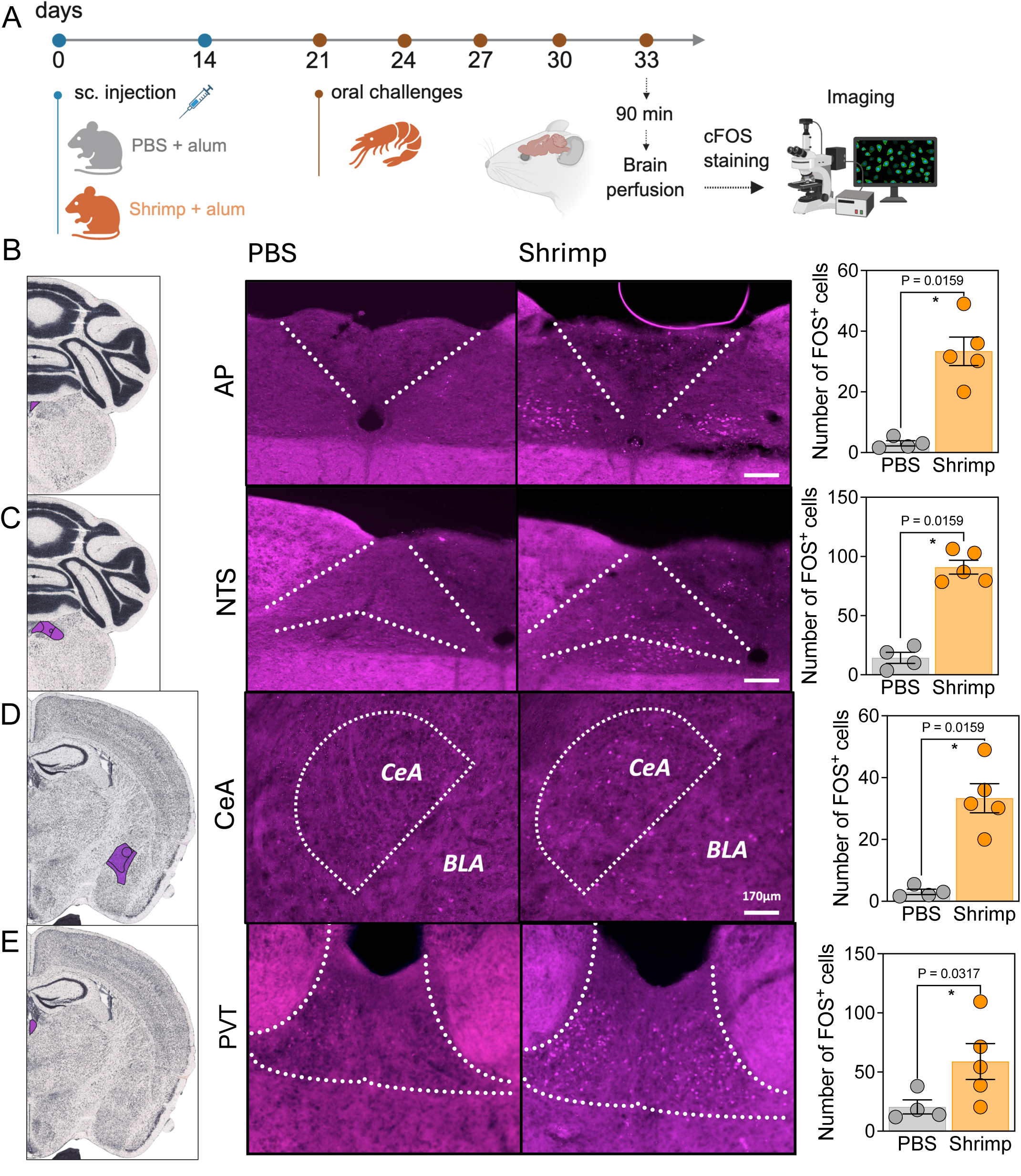
Shrimp allergy induces activation of brain areas related to aversion and sickness behaviors. (**A**) Schematic protocol for allergic sensitization, oral challenges and immunofluorescence using anti-FOS antibody, 90 min after the fifth oral challenge. (**B**) Area postrema (AP): image from brain atlas (left), representative immunofluorescence images of the area from PBS or shrimp + alum-sensitized mice (middle) and cFOS quantification (right). (**C**) Nucleus of tractus solitarius (NTS): image from brain atlas (left), representative immunofluorescence images of the area from PBS or shrimp + alum-sensitized mice (middle) and anti-cFos quantification (right). (**D**) Central amygdala (CeA): image from brain atlas (left), representative immunofluorescence images of the area from PBS or shrimp + alum-sensitized mice (middle) and cFOS quantification (right). (**E**) Paraventricular nucleus of the thalamus (PVT): image from brain atlas (left), representative immunofluorescence images of the area from PBS or shrimp + alum-sensitized mice (middle) and cFOS quantification (right). Scale bars, 170 µm. Graphs shown mean±s.e.m. \**P*≤0.05, \*\**P*≤0.01, \*\*\*\**P*≤0.0001. Two-tailed Mann– Whitney test. Each panel is representative of at least two independent experiments. (**A**) Created with BioRender.com.

Repeated oral shrimp exposure increased cFOS-positive cell counts in the area postrema (AP), NTS, CeA, and paraventricular nucleus of the thalamus (PVT) in shrimp-sensitized mice compared with controls (Figures 4B–E). In contrast, no differences in cFOS⁺ cells were found in the parabrachial nucleus (PBN), paraventricular nucleus of the hypothalamus (PVN), or hippocampus (data not shown).

These findings indicate that chronic shrimp allergy activates discrete brain regions involved in visceral sensory processing and aversive states. In particular, activation of the AP–NTS axis is consistent with detection of peripheral inflammatory or humoral signals, whereas recruitment of the CeA and PVT suggests engagement of neural circuits associated with aversion and behavioral state regulation.

### Shrimp allergy-induced avoidance depends on 5-lipoxygenase activity

We previously found that avoidance behavior in an OVA-induced food allergy model is mediated by cysteinyl leukotrienes^19,20^. The 5-lipoxygenase (5-LO) pathway is the major source of leukotrienes, potent proinflammatory lipid mediators generated from arachidonic acid metabolism. During leukotriene biosynthesis, 5-LO catalyzes the oxygenation of arachidonic acid to 5-hydroperoxyeicosatetraenoic acid (5-HPETE), followed by dehydration to leukotriene A4 (LTA4), the precursor for downstream leukotrienes. To determine whether shrimp-induced avoidance depends on this pathway, shrimp-allergic mice were treated with vehicle or Zileuton, a 5-LO inhibitor, one hour before each preference test (Figure 5A).

**Figure 5.**
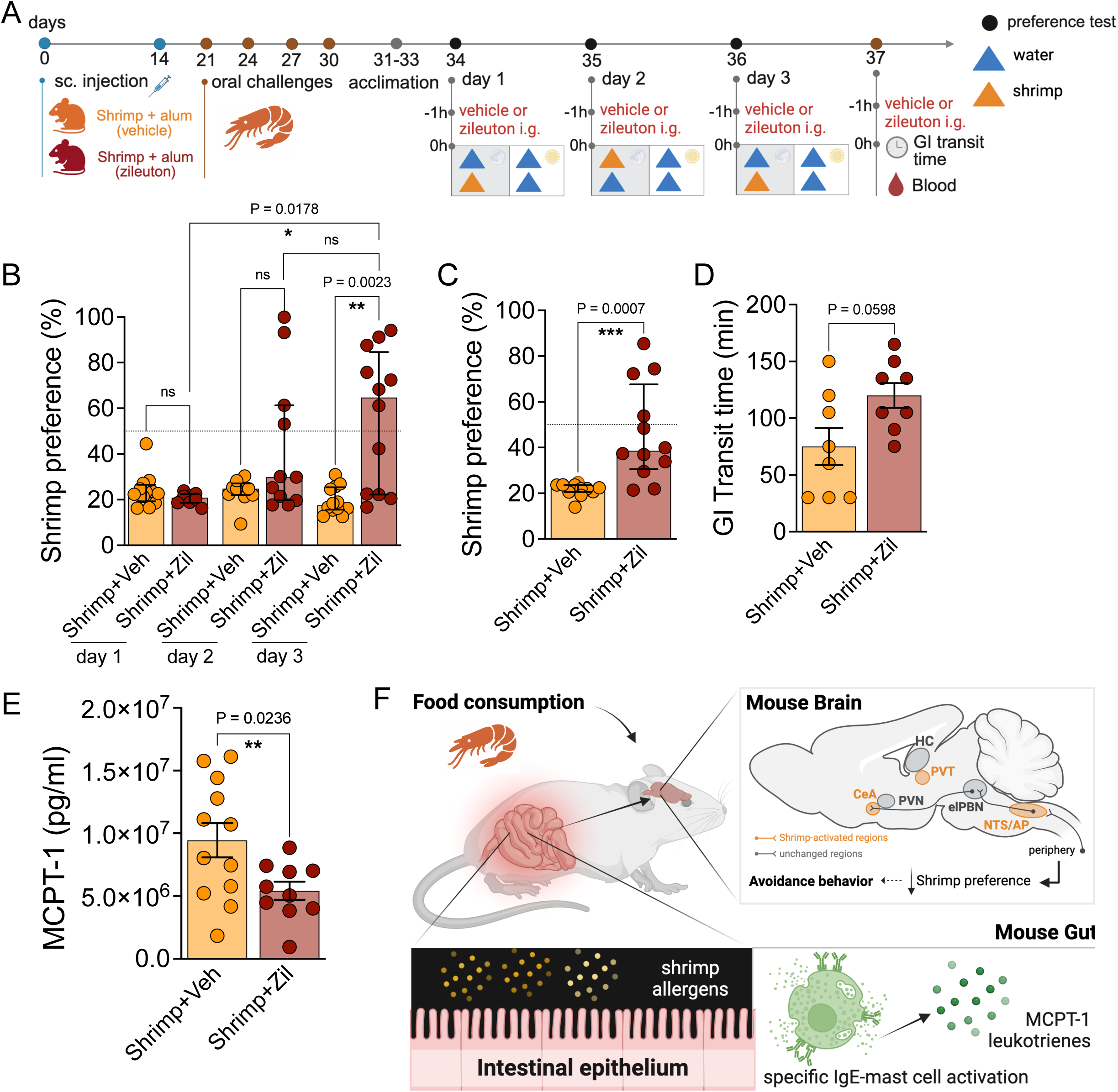
Shrimp allergy-induced avoidance behavior depends on 5-lipoxygenase activity. (**A**) Schematic protocol for allergic sensitization, oral challenges, preference test and zileuton treatment. (**B**) Shrimp preference throughout 3 days of the test with shrimp-allergic mice treated with vehicle or zileuton 1 hour before the test started (n = 12 mice/group). (**C**) Mean from shrimp preference test days with shrimp-sensitized mice treated with vehicle or zileuton 1 hour before the test started (n = 12 mice/group). (**D**) GI transit time was tracked after oral challenge (fifth) in shrimp-allergic mice treated with vehicle or zileuton 1 hour before the test started (n = 8 mice/group). (**E**) MCPT-1 serum levels from shrimp-allergic mice treated or not with zileuton 1 hour before oral challenge (n=12 mice/group). Graphs shown median ± interquartile range (**B, C**) and mean±s.e.m (**D, E**). *P≤0.05, **P≤0.01, ****P≤0.0001. Kruskal-Wallis with Dunn’s multiple comparisons test (**B**), Two-tailed Mann-Whitney test (**C, D**) and Two-tailed Unpaired T-test (**E**). Each panel is representative of one independent experiment. (**A**) Created with BioRender.com.

Zileuton treatment increased preference for shrimp-containing solution and partially reversed avoidance in allergic mice by day 3 of testing compared with vehicle-treated allergic mice (Figure 5B). Consistent with this effect, average shrimp preference across the three testing days was significantly higher in Zileuton-treated mice (Figure 5C). These results indicate that 5-LO activity contributes to shrimp allergy-induced avoidance.

We next assessed whether Zileuton altered allergic physiological responses. Given that leukotrienes have been implicated in gastrointestinal motility in OVA-induced gut inflammation^31^, we determined gastrointestinal transit after treatment (Figure 5A). Zileuton-treated allergic mice showed a trend toward altered gastrointestinal transit compared with vehicle-treated allergic mice, but this difference did not reach statistical significance (Figure 5D). In contrast, Zileuton reduced serum MCPT-1 levels compared with vehicle treatment (Figure 5E), which has also been reported in the OVA model^31^. This result suggests that 5-LO inhibition attenuates mast cell activation or its downstream consequences during repeated shrimp exposure.

Together, these findings identify 5-LO activity as a contributor to shrimp allergy-induced avoidance and mast cell activation. Overall, our data support a model in which repeated oral exposure to shrimp allergens induces IgE-associated mast cell activation and allergic inflammation in the gut, engages discrete brain regions involved in visceral sensing and aversion, and promotes leukotriene-dependent, antigen-specific avoidance behavior (Figure 5F). This response represents a selective neuroimmune defense strategy shaped by both allergic inflammation and allergen identity.

## DISCUSSION

Animals use immune, neural, endocrine, and behavioral systems to detect and respond to harmful environmental exposures. Infection-induced sickness behavior is the best-established example of this integration, but much less is known about how type 2 inflammation shapes behavior. Here, we show that repeated oral exposure to shrimp in previously sensitized mice promotes avoidance rather than a generalized sickness-like state. This behavior emerged only after repeated allergen exposure, coincided with mast cell activation and altered gastrointestinal physiology, was associated with activation of brain regions involved in visceral sensing and aversion, and depended on 5-lipoxygenase activity. These findings extend immune-mediated avoidance beyond OVA-based models and suggest that allergen avoidance is a shared, but context-dependent, neuroimmune defense strategy in food allergy.

Developing experimental models that capture clinically relevant features of shellfish allergy has been challenging because shrimp contains a complex repertoire of allergens^32,33^, and prior models have often relied on purified or recombinant proteins, strong mucosal adjuvants, or exposure routes that may not reflect how humans encounter food allergens^26,27^. Earlier tropomyosin-based models induced key features of type 2 immunity and intestinal inflammation, including allergen-specific antibodies, Th2 cytokines, eosinophil and mast cell infiltration, goblet cell hyperplasia, MCPT-1 release, and epithelial remodeling^34,35^. Our previous model using whole shrimp extract induced only a mild allergic phenotype^28^. This study sought to determine whether avoidance behavior previously described in OVA-induced food allergy also occurs in response to a more complex dietary allergen. OVA (*Gal d* 2) is a well-defined hen egg allergen and a powerful single-protein model^36^, but allergic individuals typically encounter food allergens within complex foods containing multiple allergenic and non-allergenic components^25^. Shrimp allergy therefore offers a useful system to test whether allergen-induced gut–brain communication generalizes beyond single-protein models. Using whole shrimp extract and repeated intragastric exposure, we induced key features of experimental food allergy, including elevated total IgE, shrimp-specific IgG1, MCPT-1 release, accelerated gastrointestinal transit, mild oral anaphylaxis, and intestinal remodeling. Thus, this model provides a platform to study how a complex food allergen engages immune, neural, and behavioral responses.

One of the main findings of this work is that shrimp-induced avoidance developed only after repeated oral allergen exposure. This differs from OVA-induced food allergy, in which sensitized animals can avoid the allergen after a single oral challenge^16,18–20^. In our model, shrimp-sensitized mice had elevated total IgE after sensitization but did not avoid shrimp solution until repeated challenges had induced mast cell activation, altered gastrointestinal motility, mild hypothermia, and intestinal remodeling. One possible explanation is that this protocol generates a milder allergen-specific effector response than OVA-based models, leading to weaker gut inflammation, delayed avoidance, and no detectable anxiety-like phenotype. It is possible that protocols capable of inducing stronger shrimp-specific effector responses (e.g., increased production or higher affinity IgE antibodies) would produce more robust inflammation and behavioral outputs, perhaps more closely resembling those observed in OVA-based models.

Shrimp-induced avoidance can be viewed within the broader framework of conditioned taste aversion, in which animals learn to avoid a flavor or food cue associated with an adverse internal state^37^. In this case, however, the aversive signal is generated by allergen-specific immune activation rather than by a classical toxin or pharmacological malaise-inducing stimulus^9^. Shrimp allergy therefore provides an example of immune-mediated food aversion, in which type 2 inflammation assigns negative value to a dietary cue and reduces future intake. Importantly, allergic mice avoided the shrimp-containing solution without persistent reductions in locomotor activity or consistent changes in anxiety-related or immobility measures at the time points tested. Because the OFT and EPM were performed 24 hours after allergen challenge, these findings do not exclude transient reductions in activity during the acute phase of mast cell activation and mediator release. Instead, they indicate that shrimp allergy did not produce a sustained generalized sickness-like state^38^ after the animals had recovered from the immediate reaction.

The absence of detectable anxiety-like or depressive-like behavior in our model is informative but should not be interpreted as evidence that food allergy cannot affect affective state. Clinical studies link food allergy with anxiety, reduced quality of life, and distress related to accidental exposure, but these outcomes likely reflect multiple factors, including disease severity, prior reactions, uncertainty, and learned threat expectations^10,11,39–42^. In our model, repeated shrimp exposure was sufficient to induce selective allergen avoidance, but not broader affective-like changes at the 24 h time point. Stronger allergen-specific IgE responses, longer exposure periods, repeated adverse experiences, or behavioral testing closer to the peak of allergic mediator release may be required to capture transient or sustained anxiety-related changes. In addition, OFT, EPM, and FST paradigms reflect complex behavioral dimensions beyond anxiety-like or depressive-like states and should be interpreted with caution. It is also possible that allergy-associated threat responses are context-specific and emerge primarily during feeding or in the presence of allergen-associated sensory cues rather than in standard behavioral arenas. Future experiments presenting shrimp-associated olfactory or gustatory cues during behavioral testing could help distinguish generalized anxiety-like behavior from cue-specific anticipatory threat.

Repeated shrimp exposure activated discrete brain regions involved in visceral sensation, aversion, and internal-state regulation. Activation of the AP and NTS is consistent with engagement of brainstem pathways that monitor peripheral physiological and humoral signals. The AP is positioned to detect circulating mediators and is strongly implicated in nausea, malaise, and conditioned flavor avoidance^43–45^, whereas the NTS integrates vagal and spinal sensory inputs from the gastrointestinal tract^46^. Notably, AP activation was not reported in the OVA food allergy model^18,20^, making this a potentially important distinction of shrimp-induced allergic inflammation. Because the shrimp and OVA models differ in both allergen composition and challenge history, the present study cannot determine whether AP recruitment and the absence of detectable PBN activation reflect allergen identity, repeated exposure, differences in inflammatory magnitude, or other protocol-specific factors.

Shrimp allergy also activated the CeA and PVT. CeA activation is consistent with prior work in OVA-induced allergy, where this region is recruited by allergen exposure and linked to aversive behavior^18,20,29^. The PVT finding is particularly interesting because this region integrates internal state, arousal, and prior experience to guide adaptive responses^47–49^. Its activation may relate to the progressive nature of shrimp avoidance, which emerged only after repeated allergen exposure. Together, these activation patterns (i.e., AP–NTS and CeA–PVT) suggest that shrimp allergy engages both brainstem visceral-sensing pathways and limbic-thalamic circuits involved in assigning behavioral significance to internal state. Still, cFOS mapping identifies candidate regions activated during allergen exposure but does not establish necessity or sufficiency. Future circuit-level studies will be needed to determine whether these regions are required for immune-mediated avoidance or for top-down regulatory pathways.

Our data further implicate the 5-lipoxygenase pathway in shrimp allergy-induced avoidance. 5-lipoxygenase catalyzes the conversion of arachidonic acid into leukotriene A4, the precursor of downstream leukotrienes, including cysteinyl leukotrienes. These lipid mediators are rapidly generated during IgE-dependent mast cell activation and can act on epithelial, immune, and neuronal targets^50^. In OVA-induced food allergy, cysteinyl leukotrienes are required for avoidance behavior and contribute to anaphylactic responses following oral allergen exposure^20,31^. Here, pharmacological inhibition of 5-lipoxygenase with zileuton increased consumption of shrimp-containing solution and reduced serum MCPT-1, suggesting that leukotriene-related pathways contribute to both allergic effector responses and the maintenance of avoidance during repeated shrimp exposure. Notably, the effect of zileuton on shrimp preference became progressively more apparent across the three testing days rather than occurring immediately after the first treatment. Leukotriene D_4_ (LTD4) was recently shown to enhance intestinal allergen absorption and promote oral anaphylaxis in susceptible mice^51^. The repeated 5-lipoxygenase inhibition in our model may therefore have limited allergen uptake and the ensuing allergic response during each preference test, progressively weakening the negative post-ingestive consequences associated with shrimp consumption. It is also possible that this reduction in adverse reinforcement allows updating or extinction of the previously acquired aversion when shrimp consumption is no longer followed by the same inflammatory consequences. The current design, however, cannot distinguish whether zileuton primarily affected allergen absorption, acute aversive signaling, the expression of avoidance, or relearning across successive exposures.

Several limitations should be considered. Although the use of whole shrimp extract better represents exposure to a complex food allergen, its compositional complexity will require future studies to identify the specific allergens and other food components responsible for sensitization, inflammation, neural activation, and avoidance. In addition, the absence of a validated assay for shrimp-specific IgE prevented us from directly quantifying the magnitude of the allergen-specific IgE response. Although more robust than previous shrimp allergy models, the current model does not reproduce the severe systemic anaphylaxis that can occur in shellfish-allergic patients. Because body temperature was measured at 30-minute intervals, an earlier temperature nadir may have been missed. These data therefore demonstrate mild hypothermia but do not define the acute kinetics or maximal severity of the oral allergic reaction. Finally, because our mechanistic studies relied on pharmacological inhibition of 5-lipoxygenase, future work should identify the cellular sources of the leukotrienes relevant to avoidance, including potential hematopoietic sources (e.g., mast cells) and epithelial sources (e.g., tuft cells); determine whether the behavioral response is mediated by LTB4 or cysteinyl leukotrienes; and define the relevant downstream receptors, including BLT1 and BLT2 for LTB4 and CysLT1R, CysLT2R, and OXGR1/CysLT3R, for cysteinyl leukotrienes.

Overall, this work supports a model in which repeated oral exposure to shrimp allergens induces IgE-associated mast cell activation and intestinal allergic inflammation, leading to 5-lipoxygenase-dependent engagement of gut–brain pathways that promote selective allergen avoidance. These findings position food allergy as a useful framework for studying how type 2 inflammation shapes behavior and extend immune-mediated avoidance beyond OVA-based models^21^. More broadly, our data suggest that avoidance may represent a shared neuroimmune defense strategy, whose timing, inflammatory threshold, and neural circuitry are influenced by allergen identity and exposure context.

## MATERIAL AND METHODS

### Animals

All animal care and experimentation were performed according to institutional and national guidelines for animal care and use. Animal experiments conducted in Brazil were approved by the Animal Ethics Committee on the Use of Animals of the Oswaldo Cruz Foundation – Gonçalo Moniz Institute (protocol no. 013/2023). Animal experiments conducted in the United States were approved by the Institutional Animal Care and Use Committee of Arizona State University. Female BALB/c mice (6–8 weeks old), originally from Charles River Laboratories (#000651), were used for all experiments. Mice were maintained at pathogen-free animal facilities in temperature (22°C) and humidity-controlled rooms, in a 12-hour light/dark cycle with free access to standard chow diet and water.

### Shrimp extract preparation

Whole shrimp extract was obtained following the protocol previously published by our group^28^. Briefly, total protein extract was obtained from peeled and precooked industrialized *Litopenaeus vannamei* shrimp (BrazilianFish Ltda., Brazil) using the protocol proposed by Ayuso and colleagues^52^, with some adaptations. To generate a uniform homogenate, shrimp samples were crushed and resuspended in 1X PBS solution (0.05% w/v), then incubated overnight. This suspension was first centrifuged at 1000 × *g* for 10 min at 4 °C. The supernatant was then centrifuged for 5 min at 20,000 × *g* at 4 °C. Finally, the resulting supernatant was collected and stored at –20 °C until used for experiments. For the avoidance behavior test, the shrimp homogenate was resuspended in drinking water instead of PBS, to eliminate a possible confound from preference for PBS. Total protein concentration was measured using a Pierce Micro BCA Protein Assay Kit (ThermoFisher Scientific)

### Allergic sensitization and challenges

Mice were sensitized subcutaneously on days 0 and 14 with 5 mg/kg shrimp extract adsorbed in 50 mg/kg alum gel (Invivogen vac-alu-250) and diluted to a final volume of 0.2 mL in PBS pH 7.4. Controls received all the above except for the shrimp extract (referred to as Control). For allergen challenges, all groups received 5 oral gavages with 20 mg shrimp extract in 0.25 mL of normal mice drinking water on days 21, 24, 27, 30, and 33, unless otherwise stated.

### Preference test

Drinking behavior was determined using the two-bottle preference test, based on the well-established sucrose preference test and previous studies^53,54^. Two days after booster sensitization (day 16), mice were individually housed in new cages adapted with two amber glass dropper bottles (30 mL; Seven Glass, Brazil) filled with filtered water. During a 5-day acclimation period (days 16–20), animals had continuous access to both bottles. Baseline water intake and side preference were recorded on days 19 and 20 by weighing each bottle before and after the dark phase.

On day 21, mice received one bottle containing water and one bottle containing 0.5% shrimp extract prepared in drinking water, which were placed on the side corresponding to the higher baseline water intake for each animal to minimize side bias. Bottles were introduced 1 h before lights off (ZT11) and remained in place overnight. On day 22 at ZT4, both bottles were replaced with water to reassess baseline intake during a washout period. At ZT11 on days 22 and 23, mice again received one bottle of water and one bottle of fresh 0.5% shrimp extract solution; the left–right position of the shrimp bottle was switched between days to control for side preference.

The same two-bottle preference protocol was repeated after the fourth oral shrimp challenge, performed on day 30, using the same sequence of acclimation, baseline assessment, and preference test. For each day, bottle weights were measured immediately before placement and after the test to calculate solution intake. Shrimp preference was expressed as the percentage intake of the shrimp solution relative to total intake (shrimp + water intake). Mice had ad libitum access to standard chow diet throughout the experiments.

### Behavioral assays

The open field test (OFT) was conducted 24 hours after either the first or the fifth oral challenge using cubic plastic arenas (45.2 × 45.2 × 30.4 cm)^55^ with a flat floor and matte, textured walls to avoid reflections. The elevated plus maze (EPM) was conducted 24 hours after the sixth oral challenge using a maze made of acrylic plastic with two open arms and two closed arms^56^ and elevated 100 cm from the floor. After one hour of acclimation to the behavior room, each mouse was placed in the center of the arena and monitored for 10 min using a tracking software to assess locomotion, speed, and location^57^. The arena was sanitized with 70% ethanol between each mouse. For the forced swim test (FST), mice were placed individually in water (∼25°C) for 6 min^58^. Latency to immobility and total immobility time were quantified by an investigator blinded to experimental group. Tests were recorded using a video camera (Philips SPC 611NC) connected to a computer and installed above the apparatus. Behavioral data from OFT and EPM experiments were automatically analyzed using the ANY-maze Video Tracking System software (StoeltingCo©), while the FST video scoring was conducted by an experienced researcher unaware of the experimental groups until all analyses were finalized.

### Brain preparation and immunofluorescence

Brains were collected 90 min after the fifth oral challenge with shrimp extract. To minimize gavage-induced stress, we administered sham gavages with normal drinking water for 3 days before the final allergen challenge. Mice were deeply anaesthetized with isoflurane (Covetrus) and were transcardially perfused with PBS followed by freshly prepared 4% paraformaldehyde (PFA) in PBS. Dissected brains were kept in 4% PFA at 4°C for 48 h, washed 3 times in PBS and transferred to a 30% sucrose in PBS solution for 2 days, frozen and then sliced into 40-μm-thick coronal sections (area postrema, NTS, PVT, PVN and PBN) and 100-μm-thick sections (lateral hypothalamus and CeA) using a Leica CM3050 S cryostat (Leica Biosystems). Briefly, the sections were permeabilized with PBS with 0.3% Triton X-100 for 30 min at room temperature and then blocked in PBS with 0.3% Triton X-100 and 10% normal donkey serum in 0.3 M glycine for 1 h at room temperature. Blocking was followed by incubation with rabbit monoclonal anti-FOS primary antibody (1:1,000 dilution, Cell Signaling #2250S) overnight for 16 h and then with Alexa Fluor 488-conjugated donkey anti-rabbit IgG secondary fluorescent antibody (1:500 dilution, Invitrogen A21202) for 2 h at room temperature. After being washed with the permeabilization solution again, the sections were mounted on slides and visualized by using a fluorescent All-In-One Keyence microscope (model BZ-X710, Keyence). Images were taken using the ×4 objective. Brain regions were defined based on the Allen Mouse Brain Atlas reference atlas (https://mouse.brain-map.org/) and processed using the open-source Fiji-ImageJ software. A blinded investigator manually quantified FOS+ cells throughout the entire procedure. All images were processed at the Biodesign Institute, Arizona State University.

### Antibody quantification

Serum levels of total IgE were determined by sandwich enzyme-linked immunosorbent assay (ELISA, 555248, BD Biosciences). Briefly, plates were coated with diluted (1:250) anti-mouse IgE capture antibody in carbonate buffer (pH 9.6) and incubated overnight at 4 °C. Plates were then washed three times with PBS containing 0.05% Tween-20 and blocked with 10% PBS-SBF for 1.5 h at room temperature. After three washes, serum samples were diluted 1:100 and incubated for 2 h at room temperature. Purified mouse IgE (557079, BD Biosciences) was used as the standard, with a highest concentration of 500 ng/mL followed by twofold serial dilutions to generate the standard curve. Plates were washed five times and incubated with biotinylated anti-mouse IgE detection antibody (1:500) and streptavidin-HRP (1:250) for 1 h at room temperature. In the final step, plates were washed seven times and incubated with 50 µL/well of TMB substrate solution in the dark. The reaction was stopped by adding 25 µL/well of sulfuric acid, and absorbance was measured at 450 nm using an automated ELISA reader.

Shrimp-specific IgG1 serum levels were determined by indirect ELISA. Briefly, plates were coated overnight at 4°C with 10 µg/mL of shrimp extract diluted in 0.1 M sodium carbonate buffer (pH 9.5). Plates were washed three times with PBS containing 0.05% Tween-20 between each step. Following blocking with 1% bovine serum albumin (BSA) in PBS for 2 h at room temperature, serum samples diluted 1:200 were added to the wells and incubated for 1 h at room temperature. After incubation, plates were washed three times, and 50 µL/well of HRP-conjugated anti-mouse IgG1 detection antibody (Life Technologies), diluted 1:1000, was added and incubated for 1 h at room temperature. Following three additional washes, the enzymatic reaction was developed using TMB substrate, and absorbance was measured at 450 nm using an automated ELISA reader.

### Oral anaphylaxis and gastrointestinal motility

To determine the occurrence of oral anaphylaxis, sensitized mice were challenged with 20 mg of shrimp extract intragastrically on day 33. Rectal temperature was measured every 30 min for 120 min after challenge using a probe (Thermalert TH-5). Gastrointestinal transit time was assessed immediately following the fifth oral challenge with intragastric shrimp, on day 30 after allergic sensitization at ZT4. Mice were gavaged with a 0.25 mL solution with 6% red carmine (C1022, Sigma-Aldrich), 0.5% methylcellulose (M0512, Sigma-Aldrich), and 20 mg shrimp extract. After oral gavage and for the duration of the assay, mice were individually placed in clean cages containing normal bedding. Mice had free access to food and water and were monitored for the occurrence of diarrhea. The gastrointestinal transit time was measured as the time between oral gavage and the appearance of the first fecal pellet containing the red carmine dye. Mice were grouped at the end of the assay.

### MCPT-1 quantification

Serum for all MCPT-1 measurements was taken from mice 1 h after 20 mg intragastric shrimp challenge unless otherwise noted. MCPT-1 levels in the serum were measured by ELISA (88-7503-22, Invitrogen).

### 5-lipoxygenase inhibition

Zileuton 50 mg/kg (Z4277, Sigma) was administered 1 hour before each day of the preference test or starting 1 h before the fifth oral challenge following GI transit time evaluation. Zileuton was administered via oral gavage in sterile 0.6% methylcellulose. The control group received only the vehicle solution.

## Statistical analysis

Data normality was assessed via D’Agostino-Pearson. Parametric data (ANOVA + Tukey’s) and non-parametric data (Kruskal-Wallis + Dunn’s) were analyzed. Results are presented as means ± standard deviation (SD) or medians and interquartile range (IQR). P values < 0.05 were assigned as statistically significant. All statistical analyses were performed using GraphPad Prism v.11.0.1 software (San Diego, CA, USA).

## Author Contributions

CMA: Conceptualization, Methodology, Formal analysis, Data curation, Visualization, Writing – original draft, Writing – review and editing. BGCL: Methodology, Formal analysis, Writing – review and editing. NSP, ACR, RCS, ISG, LMP, AMLCL, and GDS: Investigation. TUM and ACMF: Conceptualization, Supervision, Writing – review and editing. CIB: Conceptualization, Methodology, Supervision, Project administration, Writing – original draft, Writing – review and editing. EBF: Conceptualization, Methodology, Supervision, Project administration, Writing – original draft, Writing – review and editing. All authors contributed to the article and approved the submitted version.

## Acknowledgments

We thank the members of the Florsheim, Brodskyn, and Caetano Faria laboratories for helpful discussions, and the animal facility staff at both institutions for excellent animal care and husbandry. We also thank Nathaniel D. Bachtel at Yale School of Medicine for his constructive feedback, critical review of the manuscript, and valuable suggestions. This study was supported by the Instituto Nacional de Ciência e Tecnologia de Investigação em Imunologia (INCT-iii) and the INCT Mucosa e Pele. CMA’s research fellowship in the Florsheim laboratory at Arizona State University was supported in part by the Programa de Doutorado-Sanduíche no Exterior (PDSE) of the Coordenação de Aperfeiçoamento de Pessoal de Nível Superior (CAPES), Brazil. CAPES also provided fellowships to CMA, BGCL, AMLCL, and GDS (Finance Code 001), and the Oswaldo Cruz Foundation provided fellowships to IDG and NSP. NMT, TUM, AMCF, and CIB are senior investigators of the Conselho Nacional de Desenvolvimento Científico e Tecnológico (CNPq). EBF is supported by the Food Allergy Science Initiative, the Hypothesis Fund, and the Arizona Department of Health.

## Declaration of Generative AI and AI-assisted technologies

During the preparation of this work, the author(s) used ChatGPT (OpenAI) and Claude (Anthropic) to improve the clarity, conciseness, organization, and language of the manuscript. After using these tools, the authors reviewed, verified, and edited the content as needed and take full responsibility for the content of the published article.

**Supplementary Figure 1 Shrimp allergy does not produce a consistent affective-like or locomotor phenotype at the time points tested**. (**A, G**) Schematic protocol for oral challenge and open field test (OFT). (**B**) Distance traveled, (**C**) time in the center, (**D**) time in the corners, (**E**) entries in the center and (**F**) entries in the corners from OFT performed 24 hours after first oral challenge in mice sensitized with PBS or shrimp + alum (n= 15 mice per group). (**H**) Time in the center and (**I**) time in the corners from OFT performed 24 hours after the fifth oral challenge in mice sensitized with PBS or shrimp + alum (n= 15 mice per group). (**L**) Schematic protocol for oral challenge and elevated plus maze (EPM). (**M**) Time in the closed arms and (**N**) time in the open arms from EPM performed 24 hours after sixth oral challenge in mice sensitized with PBS or shrimp + alum (n=15 mice per group). Graphs shown mean±s.e.m.\*\**P*≤0.01. Two-tailed Unpaired T-test. Each panel is representative of at least two independent experiments. (**A**, **G**, **L**) Created with BioRender.com.

## Notes

### Competing Interest Statement

The authors have declared no competing interest.

