## Supplemental Figure 1 for "Repeated shrimp allergen exposure drives 5-lipoxygenase-dependent avoidance and selective gut-brain activation"

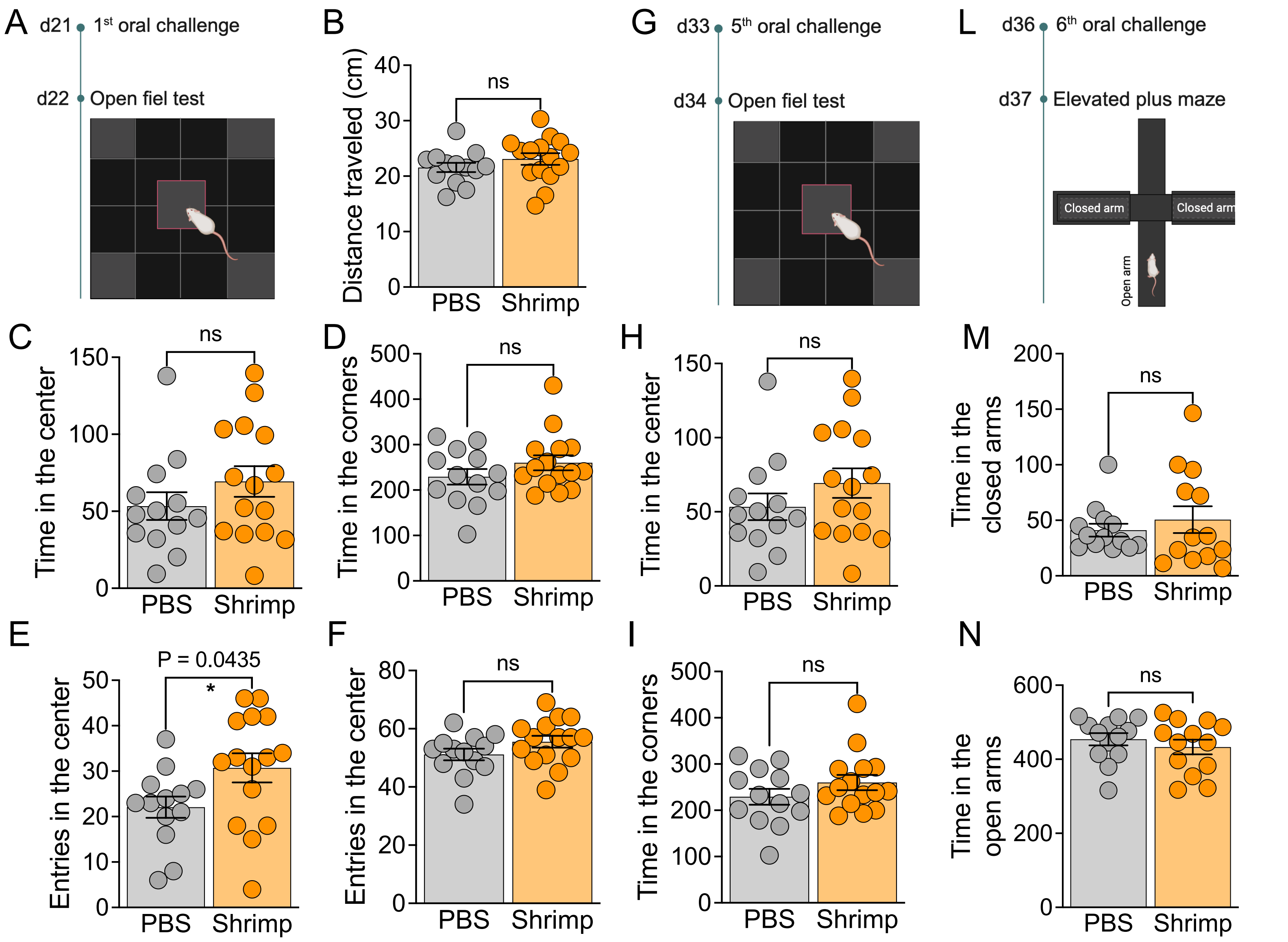


**Supplementary Figure 1 | Shrimp allergy does not produce a consistent affective-like or locomotor phenotype at the time points tested**. (**A, G**) Schematic protocol for oral challenge and open field test (OFT). (**B**) Distance traveled, (**C**) time in the center, (**D**) time in the corners, (**E**) entries in the center and (**F**) entries in the corners from OFT performed 24 hours after first oral challenge in mice sensitized with PBS or shrimp + alum (n= 15 mice per group). (**H**) Time in the center and (**I**) time in the corners from OFT performed 24 hours after the fifth oral challenge in mice sensitized with PBS or shrimp + alum (n= 15 mice per group). (**L**) Schematic protocol for oral challenge and elevated plus maze (EPM). (**M**) Time in the closed arms and (**N**) time in the open arms from EPM performed 24 hours after sixth oral challenge in mice sensitized with PBS or shrimp + alum (n= 15 mice per group). Graphs shown mean ± s.e.m.***P* ≤ 0.01. Two-tailed Unpaired T-test. Each panel is representative of at least two independent experiments. (**A**, **G**, **L**) Created with BioRender.com.
